# Infection of equine bronchial epithelial cells with a SARS-CoV-2 pseudovirus

**DOI:** 10.1101/2023.01.26.525770

**Authors:** Rebecca M. Legere, Angelica R. Allegro, Yvonne Affram, Bibiana Petri da Silveira, Jennifer L. Fridley, Kelsey M. Wells, Numan Oezguen, Robert C. Burghardt, Gus A. Wright, Jeroen Pollet, Angela I. Bordin, Paul de Figueiredo, Julian L. Leibowitz, Noah D. Cohen

**Author notes:** Address correspondence to Noah D. Cohen.

## Abstract

Severe acute respiratory syndrome coronavirus 2 (SARS-CoV-2), the causal agent of COVID-19, can infect animals by binding to the angiotensin-converting enzyme 2 (ACE2). Equine infection appears possible due to high homology (≈97%) between human and equine ACE2, evidence of *in vitro* infection in cell lines expressing equine ACE2, and evidence of seroconversion in horses after exposure to persons infected with SARS-CoV-2. Our objective was to examine susceptibility of cultured primary equine bronchial epithelial cells (EBECs) to a SARS-CoV-2 pseudovirus relative to human bronchial epithelial cells (HBECs; positive control). ACE2 expression in EBECs detected by immunofluorescence, western immunoblotting, and flow cytometry was lower in EBECs than in HBECs. EBECs were transduced with a lentivirus pseudotyped with the SARS-CoV-2 spike protein that binds to ACE2 and expresses the enhanced green fluorescent protein (eGFP) as a reporter. Cells were co-cultivated with the pseudovirus at a multiplicity of infection of 0.1 for 6 hours, washed, and maintained in media. After 96 hours, eGFP expression in EBECs was demonstrated by fluorescence microscopy, and mean Δ Ct values from quantitative PCR were significantly (P < 0.0001) higher in HBECs (8.78) than HBECs (3.24) indicating lower infectivity in EBECs. Equine respiratory tract cells were susceptible to infection with a SARS-CoV-2 pseudovirus. Lower replication efficiency in EBECs suggests that horses are unlikely to be an important zoonotic host of SARS-CoV-2, but viral mutations could render some strains more infectious to horses. Serological and virological monitoring of horses in contact with persons shedding SARS-CoV-2 is warranted.

**IMPORTANCE:** This study provides the first published evidence for SARS-CoV-2 pseudovirus infection in equine airway epithelial cells, which were less susceptible to infection than cells of human origin. This was presumably due to lower ACE2 expression in equine cells, lower viral affinity for equine ACE2, or both. Our results are important considering recent evidence for asymptomatic seroconversion in horses following exposure to COVID-19 positive humans, despite this lower susceptibility, and increased affinity of viral variants of concern for equine ACE2 compared to ancestral strains. Thus, there is great need to better characterize SARS-CoV-2 susceptibility in horses for the benefit of veterinary and human health.

## INTRODUCTION

The severe acute respiratory syndrome coronavirus 2 (SARS-CoV-2) has caused a global pandemic of the disease known as coronavirus disease 2019 (COVID-19). A wide range of species of domestic animals and wildlife have been infected by SARS-CoV-2 (1-4). Humans who handle horses have considerable direct contact with horses, including proximity to mucosal surfaces of the mouth and nose during activities such as haltering, placing bits, and feeding. Serological evidence of exposure to SARS-CoV-2 in horses has been documented after exposure to infected humans (5, 6), but clinical signs of infection in horses have not been reported to date.

Infection with SARS-CoV-2 primarily occurs by binding of the virus to the angiotensin converting enzyme 2 (ACE2). The equine and human ACE2 receptors (eACE2 and huACE2, respectively) have high homology (estimated to be 96.8%; **Fig. 1**), including the domain which binds the SARS-CoV-2 spike protein (7-9). Evidence of infectivity of SARS-CoV-2 for horses is conflicting. Virus could neither be isolated nor detected by reverse transcription PCR (RT-PCR) from nasal swabs, rectal swabs, or various tissues collected from a horse after intranasal infection with SARS-CoV-2 (10). However, SARS-CoV-2 can infect equine dermal fibroblasts and human cells expressing eACE2 *in vitro*, although to a lesser extent than cell lines expressing huACE2 (8, 11-13). To the authors’ knowledge, infection of equine respiratory tract cells with SARS-CoV-2 or a SARS-CoV-2 pseudovirus has not been reported. In this study, we examined expression of ACE2 by equine bronchial epithelial cells (EBECs) and infection of EBECs with a lentivirus construct expressing the spike protein of SARS-CoV-2 that binds to ACE2.

**Figure 1.**
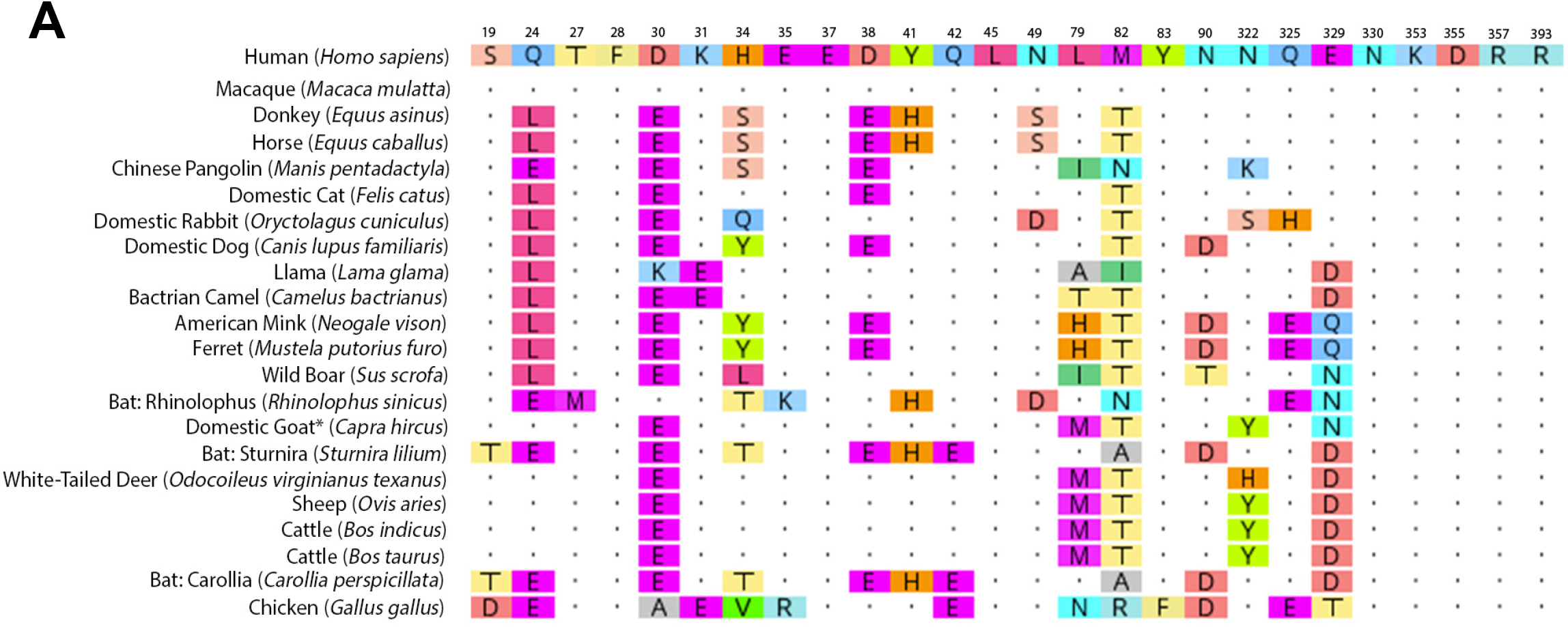
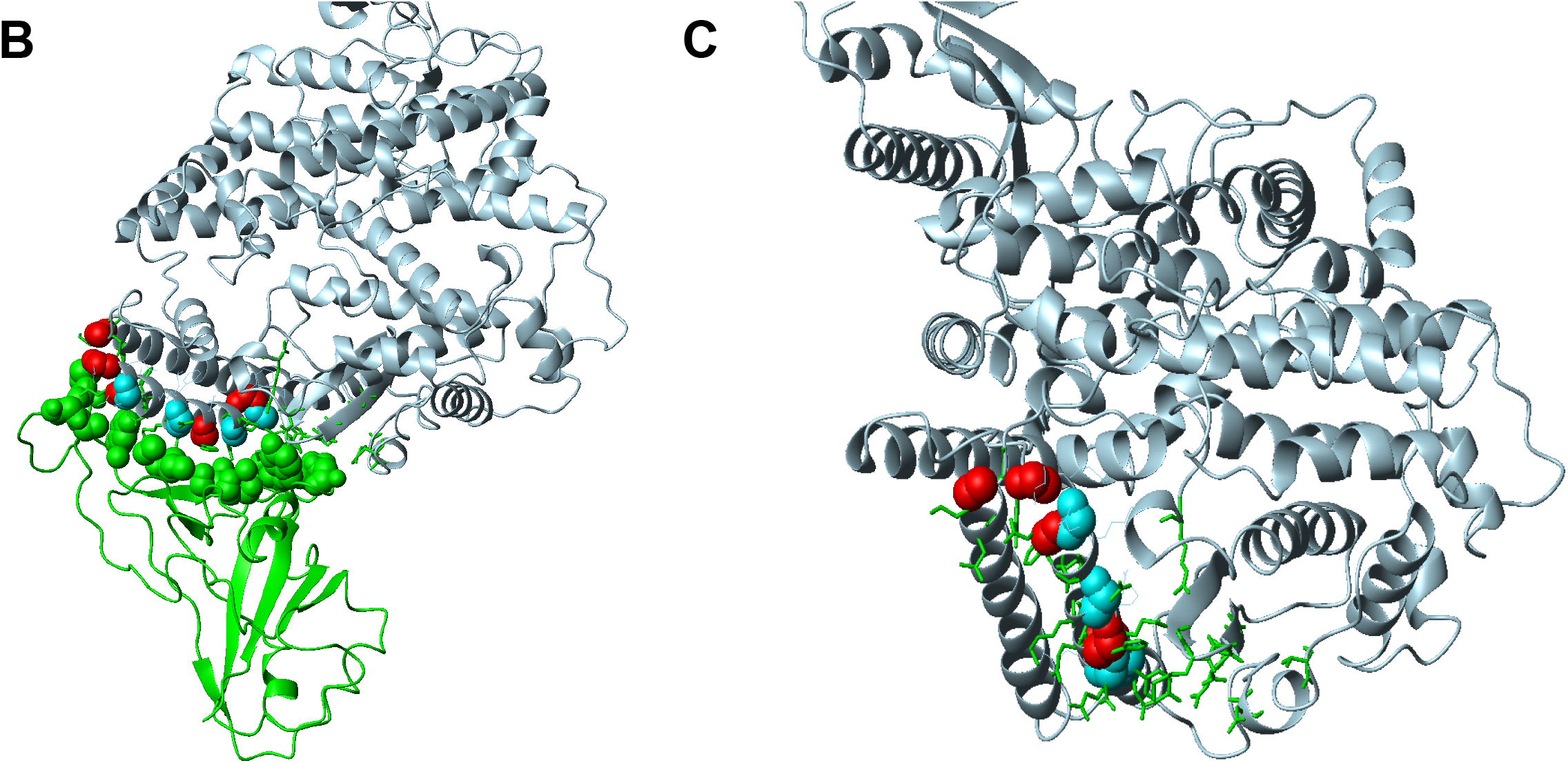
A. ACE2 Sequence Alignment B,C. 3D Equine ACE2 Protein Structure. Angiotensin-converting enzyme 2 (ACE2) protein sequences for various species and horses, and their similarity human ACE2 (huACE2). **(A)** ACE2 protein sequence alignment of species based on likely human contact and exposure (*i*.*e*., pets and livestock) or wild animals documented as susceptible to infection (*e*.*g*., bats and whitetail deer), using a scheme adapted from a previous report (14) and updated to include horses (accession data in Supplementary Table 1). Overall sequence homology to huACE2, and specific residues identified as critical for interaction between the huACE2 protein and the receptor binding domain of SARS-CoV-2 (methods adapted from the work of M. Gultom and co-authors (29)), were compared and ranked vertically by overall % homology of the ACE2 sequence to huACE2, amino acid changes are denoted by colored blocks, and ñ denotes a predicted sequence. We then generated a 3-dimensional protein structure of eACE2 at its predicted interaction site with SARS-CoV-2 spike protein receptor binding domain (RBD), and compared its similarity to huACE2 in 2 orientations **(B, C)**. Overall, sequence homology between eACE2 and huACE2 was 96.8%, and 3-dimensional structures were nearly identical between equine and huACE. Alignment between the spike protein’s receptor binding domain (RBD) and ACE2 interface revealed that this critical interface was highly similar; green regions were the portions of this critical interaction site’s sequence that were conserved between human and horse, cyan regions were homologous, and only the red regions differed between horses and humans.

## RESULTS

### ACE2 Expression

To characterize the equine ACE2 sequence *in silico*, the ACE2 protein sequences for various species, selected based on likely human contact and exposure (i.e., pets and livestock) or wild animals documented as susceptible to infection (e.g., bats and whitetail deer), using a scheme adapted from a previous report (14) and updated to include horses, were retrieved from the National Center for Biotechnology Information Protein Database (NCBI, accession data in Supplementary Table 1). When these sequences were aligned for overall sequence homology to huACE2 and compared at specific residues identified as critical for interaction between the huACE2 protein and the receptor binding domain of SARS-CoV-2 (methods adapted from the work of Gultom and collaborators (14)), horses had greater overall homology than species known to be viral reservoirs (e.g., pangolin or cats) (**Fig. 1A**), although amino acids at critical residues varied among species. Three-dimensional modelling of the equine ACE2 protein sequence (**Fig. 1B, C**) using the SWISS-MODEL web service revealed overall sequence homology between eACE2 and huACE2 was 96.8%, and that the alignment at the crucial interface between the spike protein’s receptor binding domain (RBD) and ACE2 interface was highly similar.

To demonstrate ACE2 expression in equine airway tissues, primary EBEC cultures were established according to previously described methods (15, 16), and human bronchial epithelial cells (HBECs, #C-12640, Promocell) were cultured in parallel as a positive control. Expression of ACE2 was detected in EBECs by western immunoblot (**Fig. 2**), with strongest bands detected in EBECs for the 90-kDa isotype, weakly at the 130-kDa isotype (strongest in human lung tissues), and not detected at the 55-kDa isotype. ACE2 expression was also detected in EBECs by fluorescence immunocytochemistry (**Fig. 3**) following confirmation of epithelial cell type by cytokeratin staining (**Fig. 3A, 3C**). Flow cytometry of ACE2 indicated significantly (P<0.0001) fewer EBECs (13.4%) were positive for ACE2 compared to HBECs (35.3%) (**Fig. 4**).

**Figure 2.**
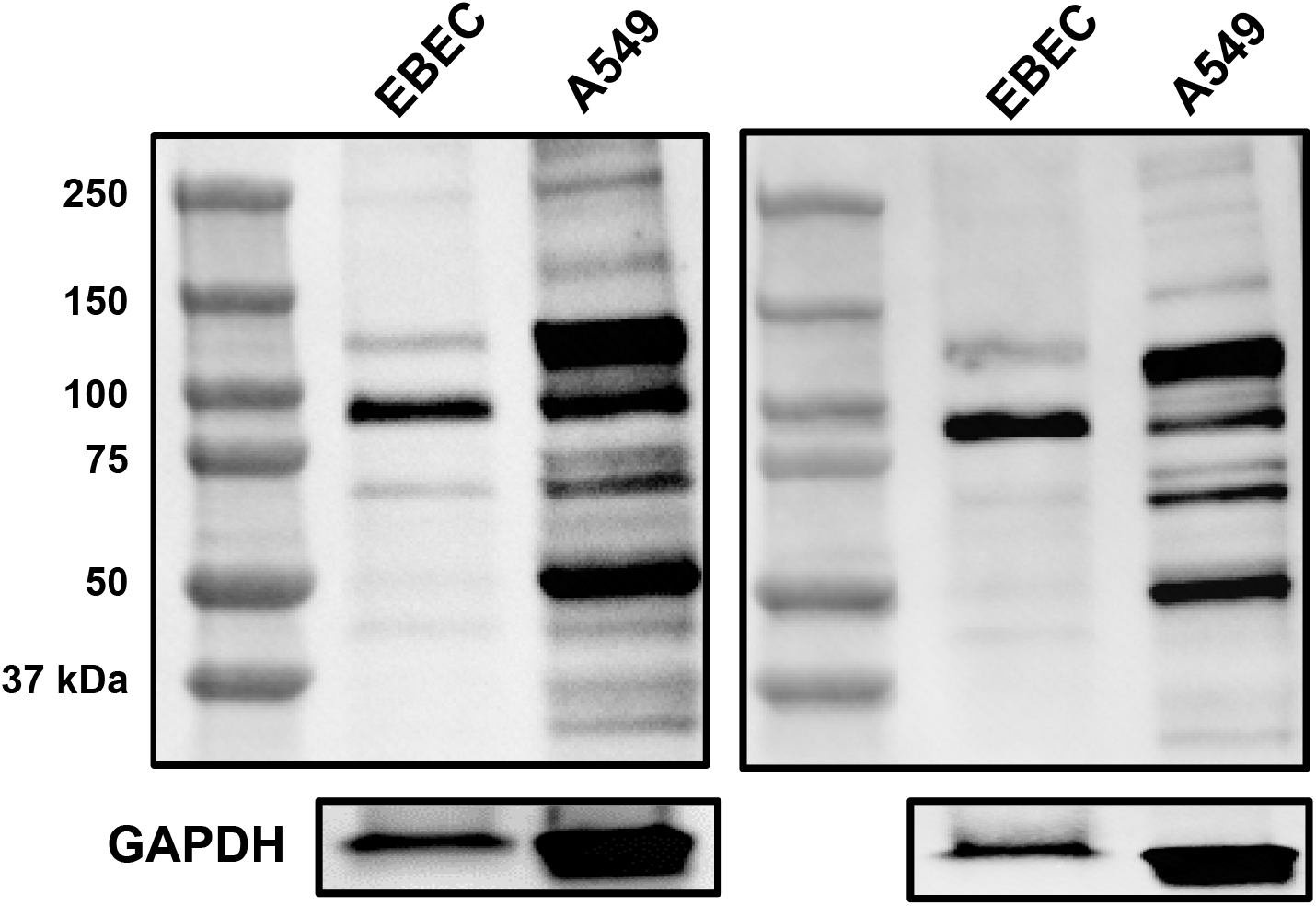
Western immunoblot for ACE2 expression in equine bronchial epithelial cells (EBECs). Lysates of EBECs and human lung adenocarcinoma cells (A549, positive control) were probed with rabbit anti-huACE2 antibody targeting the C-terminus. Bands were strongest in EBECs for the 90-kDa isotype, weakly for the 130-kDa isotype (strongest in human lung tissues), and not detected at the 55-kDa isotype. Experiments were performed with EBECs from 2 unrelated horses. An additional aliquot of each sample was probed concurrently with mouse anti-human GAPDH antibody as loading control.

**Figure 3.**
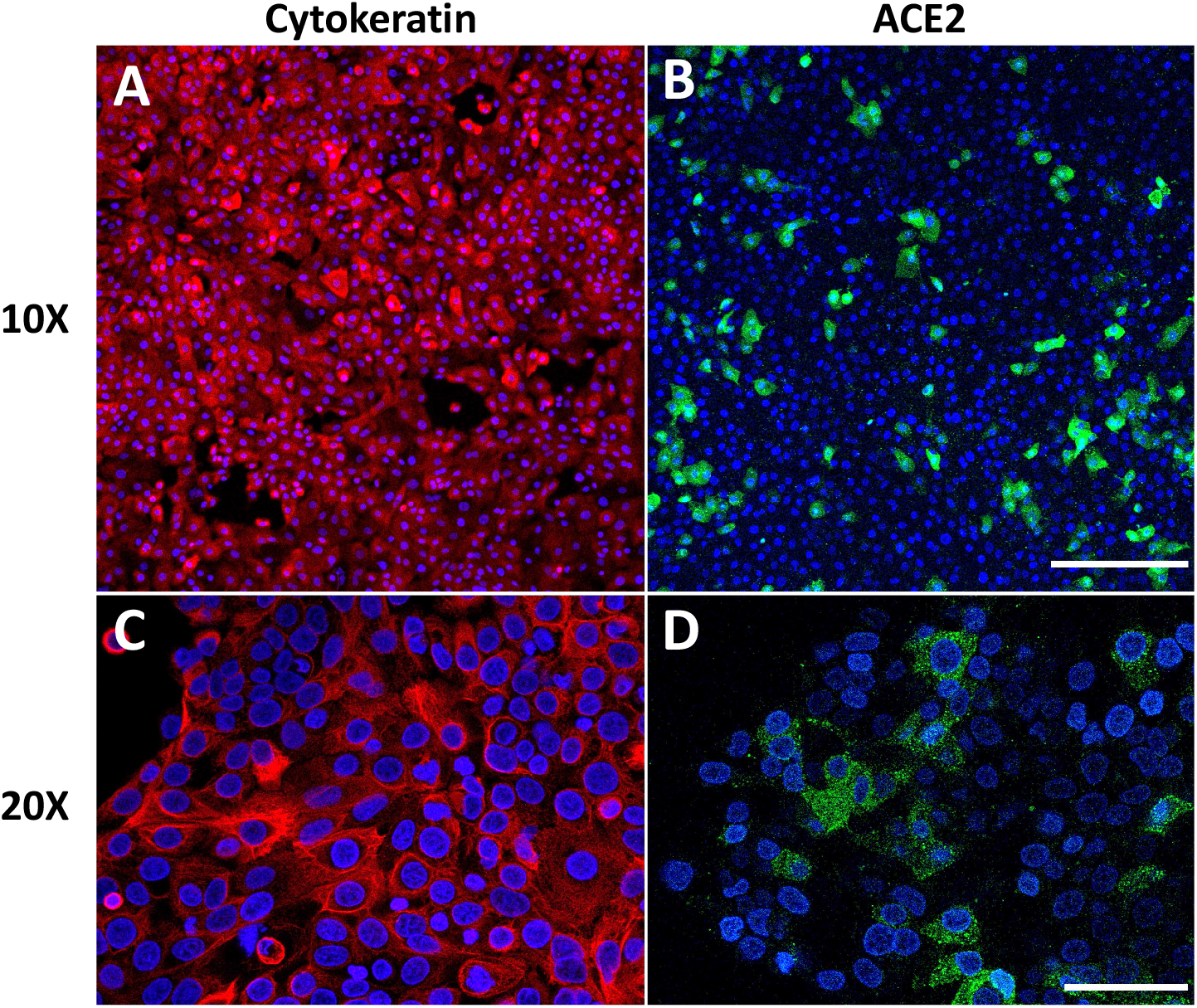
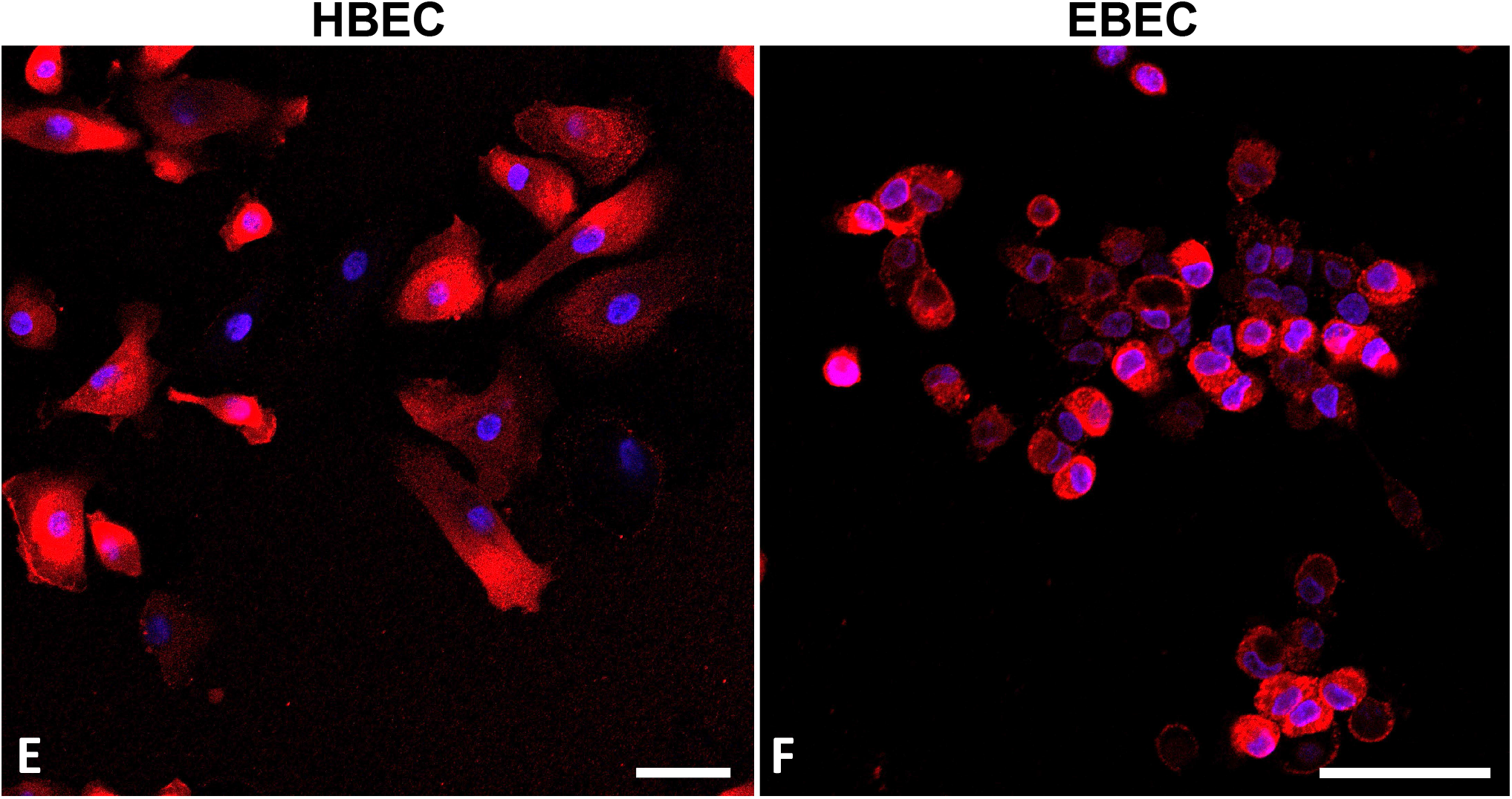
Fluorescent immunocytochemistry for expression of ACE2 and cytokeratin in equine bronchial epithelial cells (EBECs) and human bronchial epithelial cells (HBECs). Permeabilized EBECs were stained to confirm epithelial cell type **(A, C)** with monoclonal murine anti-cytokeratin AE1/AE3 antibody. Non-permeabilized EBECs were stained for ACE2 expression **(B, D)** with rabbit anti-huACE2 antibody. The top 2 pictures were imaged with a 10X lens (line scale = 200 µm); bottom 2 pictures were imaged with 20X lens (line scale = 50 µm). In a subsequent experiment, EBECs and HBECs were stained for ACE2 **(E, F)** with huACE2 antibody. Both were imaged with 40X lens, but EBECs were shown at higher magnification **(F)** than HBECs **(E)** to demonstrate staining detail (line scale = 50 µm in both images).

**Figure 4.**
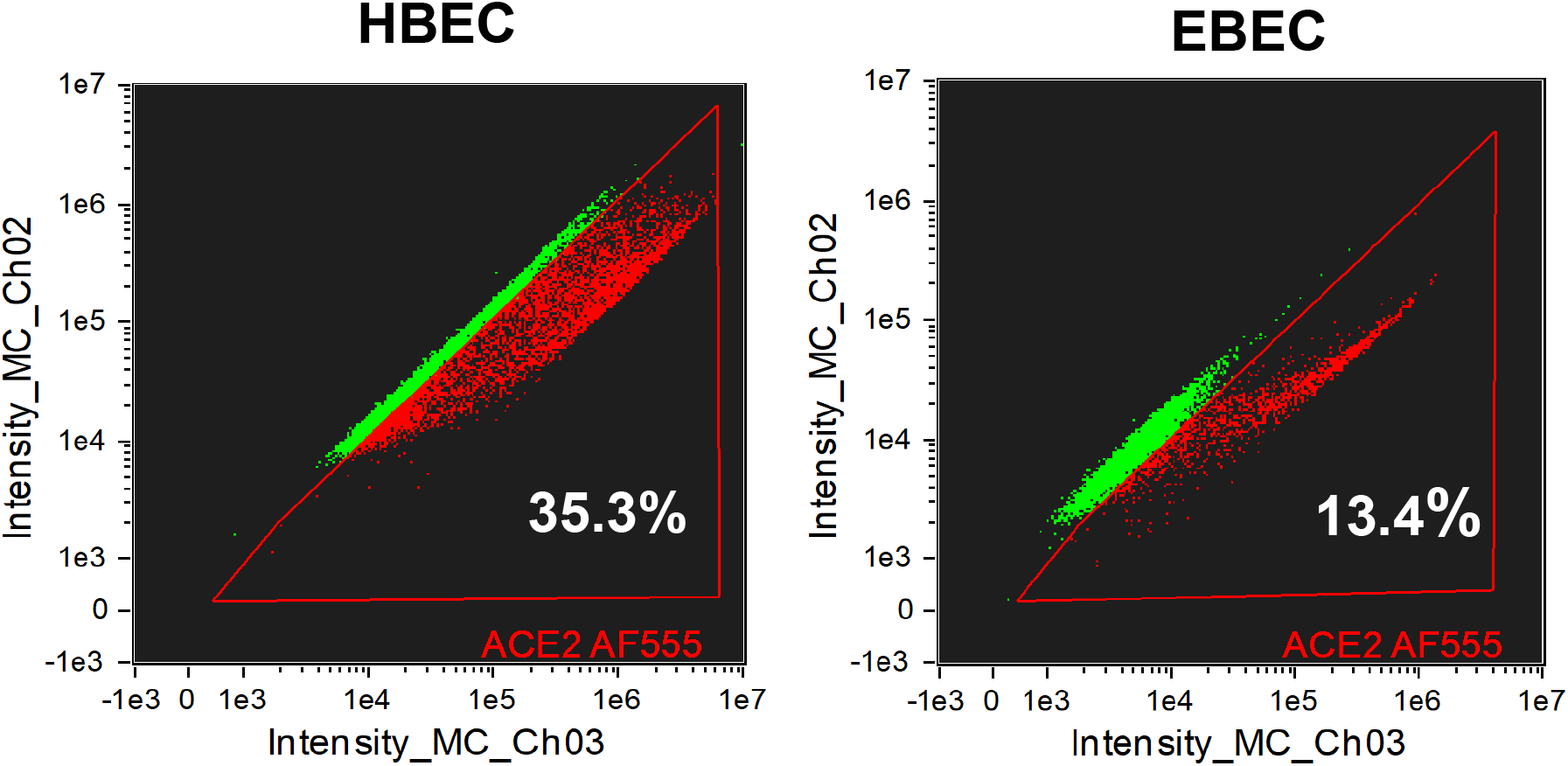
Flow Cytometry for ACE2. Imaging flow cytometry of ACE2 expression in equine bronchial epithelial cells (EBECs) and human bronchial epithelial cells (HBECs). Cultured EBECs and HBECs were fixed and stained with an affinity-purified polyclonal goat anti-huACE2 antibody targeting the extracellular domain. Gating of ACE2 positive cells indicated significantly (P<0.0001) fewer ACE-2 positive cells in EBECs (13.4%) than in HBECs (35.3%).

### Pseudovirus Transduction

To demonstrate susceptibility to SARS-CoV-2, the primary cultures of EBECs and HBECs were treated with a lentivirus pseudotyped with the SARS-CoV-2 spike protein and expressing enhanced green fluorescent protein (eGFP) as a reporter at an MOI of 0.1 for 6 hours. After 96 hours, fluorescence microscopy demonstrated green fluorescence in EBECs and HBECs transduced with the pseudovirus construct encoding *egfp* (**Fig. 5**); no green fluorescence was observed in non-transduced cells. To compare susceptibility between equine and human cells, qPCR was performed to quantify the eGFP expression within transduced cells, relative to expression of a housekeeping gene (β-2-microglobulin). eGFP Ct values were significantly (P < 0.0001) lower in transduced than non-transduced EBECs and HBECs, indicating significantly higher expression of *egfp* in transduced cells (**Fig. 6A**). The ΔCt (*i*.*e*., Ct[*egfp*] – Ct[*B2M*]) values were significantly smaller for HBECs (mean, 3.24; SD, 0.11) than EBECs (mean, 8.78; SD, 0.88) (**Fig. 6B**) indicating greater relative expression - and thus greater transduction - of *egpf* in HBECs than EBECs.

**Figure 5.**
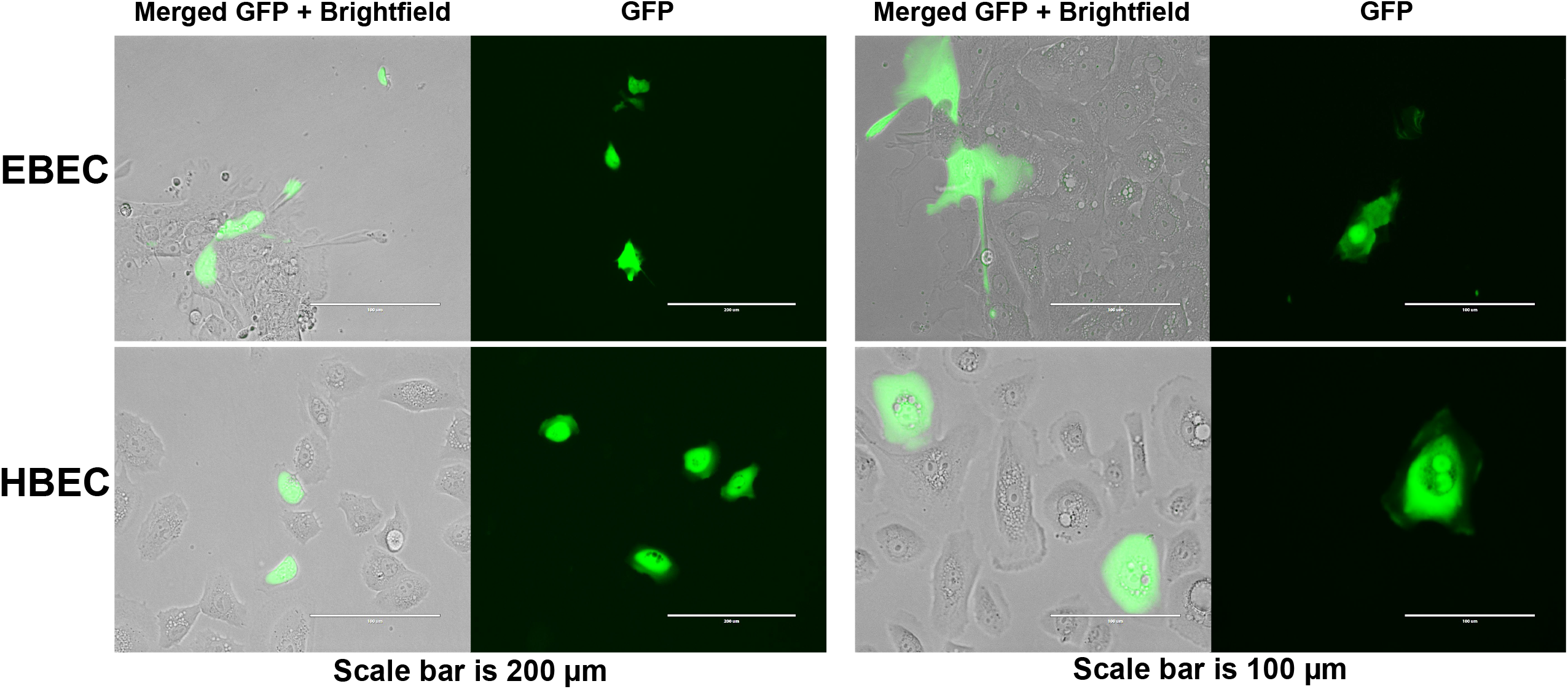
Fluorescent Microscopy. eGFP expression of equine bronchial epithelial cells (EBECs) and human bronchial epithelial cells (HBECs) imaged by fluorescent microscopy, following transduction with lentivirus pseudotyped with SARS-CoV-2 spike protein and expressing eGFP as a reporter. Cultured cells were transduced and imaged after 96 hours in brightfield and GFP channels. Representative images from experiments successfully repeated in 2 unrelated horses.

**Figure 6.**
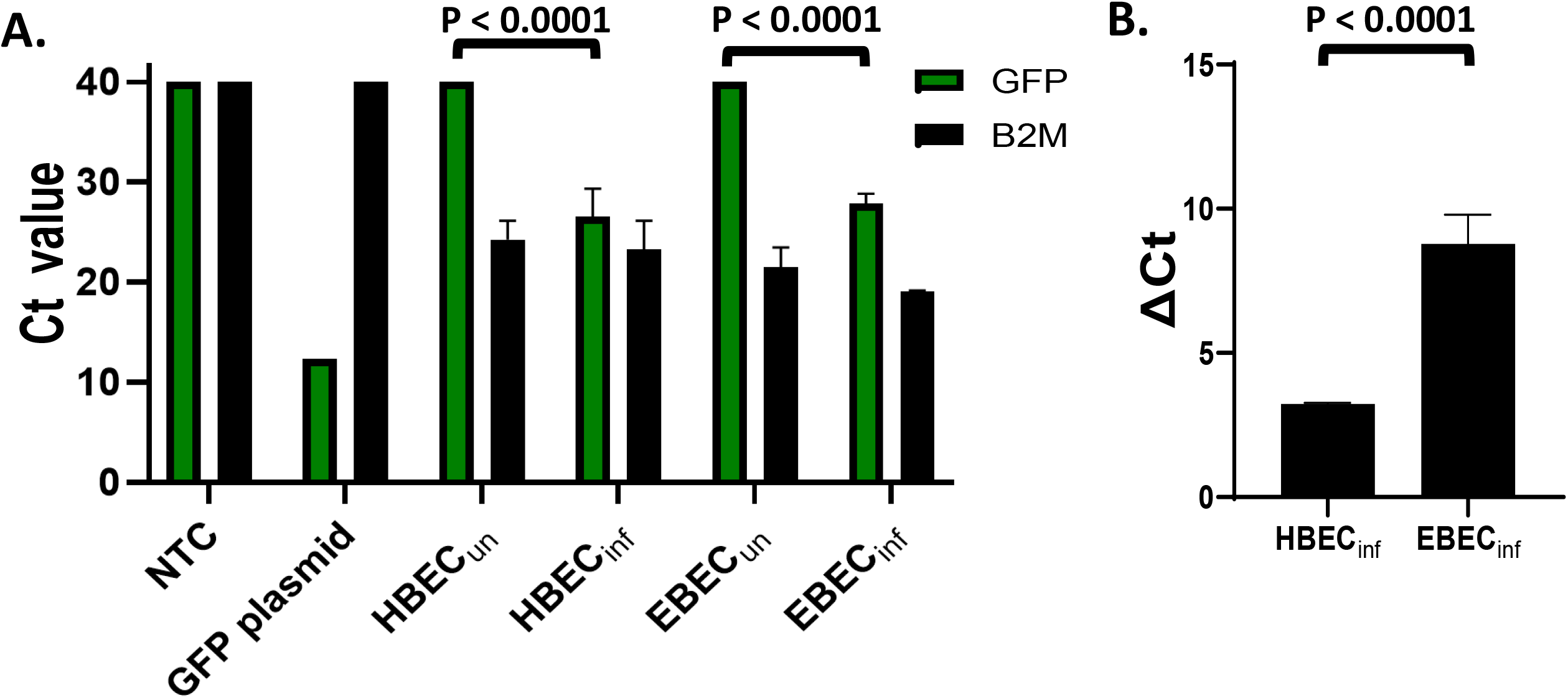
qPCR. Quantitative PCR for expression of GFP and β-2-microglobulin (*B2M*, housekeeping gene) in EBECs and HBECs, compared to a non-template control (*NTC*) and a DNA plasmid expressing GFP as positive control. Each condition was tested in triplicate, and each sample also was tested in triplicate. (**A)** Lower Ct values indicate higher GFP expression in infected wells compared to uninfected wells, but no difference between EBECs and HBECs. (**B)** The ΔCt values (Ct[***GFP***] – Ct[***B2M***]) indicate higher GFP expression in infected HBEC cells compared to infected EBECs. Error bars indicate standard deviations.

## DISCUSSION

Our findings indicate that EBECs can be transduced with a pseudovirus expressing the SARS-CoV-2 S protein, suggesting that horses can be infected with SARS-CoV-2. These findings are consistent with recent reports of seroconversion to SARS-CoV-2 in horses (5, 6). Our results also concur with *in vitro* evidence of SARS-CoV-2 infection in cells expressing eACE2, both in human cell lines transfected to express the protein (8, 11, 13) and in an equine dermal fibroblast cell line expressing ACE2 (12). Our results are plausible given the high homology between the eACE2 and huACE2, and similarity in the critical regions of the interaction site between the RBD of the S protein and eACE2 (**Fig. 1**) (9, 11, 17). Using a primary culture of EBECs isolated from healthy horses to represent a site of primary aerosol exposure to inhaled viruses in horses, we demonstrated susceptibility to infection with pseudovirus in all 3 horses tested.

The lower efficiency of transduction for EBECs than HBECs indicates that horses might be less susceptible to infection than humans. Our results regarding the relative efficiency of infection of EBECs and HBECs should be interpreted with caution because of the small number of horses tested and differences between EBECs harvested from horses and commercially-sourced HBECs. However, our findings of lower infectivity are consistent with reports suggesting that horses are not highly susceptible to infection with SARS-CoV-2. Experimental infection with SARS-CoV-2 was unsuccessful in a single horse (10) and virus could not be detected in horses with exposure to persons testing positive for SARS-CoV-2 by rt-PCR (18) or serological testing (5). Lower susceptibility of horses to infection with SARS-CoV-2 was predicted by crystallographic modelling of eACE2 (19) and was reported for non-respiratory cell lines expressing eACE2. The probability of SARS-CoV-2 entry via eACE2 has been predicted using crystallography modelling but reported estimates have ranged from >90% (20) to 48%; the latter value was comparable to that estimated for mice (*viz*., predicted entry probability of 50%), a species not used as a research model because they are considered to be at low-risk of infection with SARS-CoV-2 (19). In another study, binding affinity between eACE2 and the RBD of the S protein of SARS-CoV-2 was ∼32 times lower than that of huACE2 (9). Notably, however, this difference was halved by a single amino acid substitution in eACE2; this finding highlights the potential for some viral variants with altered binding affinity to be more infectious in horses. Human cell lines expressing eACE2 have been demonstrated to be less susceptible to infection with SARS-CoV-2 than cells expressing huACE2 (8, 11, 13). An equine dermal fibroblast cell line was successfully infected *in vitro* with SARS-CoV-2 with lower efficiency than human cell lines (12); however, dermal fibroblasts may not be representative of the respiratory epithelium (the *in vivo* target tissue for aerosol infection). Several SARS-CoV-2 pseudovirus constructs expressing the S protein of SARS-CoV-2 variants of concern were able to infect cells expressing eACE2 more readily than that of pseudovirus with the wild-type S protein, and efficiency of infection of the pseudovirus of the delta strain was equivalent to that in huACE2-expressing cells (8). Thus, it is possible that variants of SARS-CoV-2 might emerge that can more readily infect horses.

Our study has several limitations. We used a pseudovirus to demonstrate infectivity rather than SARS-CoV-2 virus. The extent to which pseudovirus infection represents infection with strains of SARS-CoV-2 and whether virus can persist within infected EBECs remain unknown. The use of a pseudovirus had the advantage of enabling us to work at biosafety level 2 conditions to demonstrate potential for infection of equine respiratory epithelial cells expressing ACE2. This pseudovirus reporter system has been used as a corollary of infection in cells expressing eACE2 (13) and is a standard assay for rapidly evaluating efficiency of cell entry and neutralization with SARS-CoV-2 for vaccine development (21, 22). The concordance of our findings of lower efficiency of infectivity of HBECS than EBECs with the pseudovirus with results of lower infectivity of SARS-CoV-2 of cells expressing eACE2 than hACE2 indicates that the pseudovirus is a valid surrogate for SARS-CoV-2. We detected ACE2 in equine cells using antibodies against huACE2. This might have lowered sensitivity for detecting eACE2 in equine tissues. Our cell culture model was maintained in submerged conditions, leading to partial differentiation and fewer ciliated cells than what would be achieved either by an air-liquid interface culture system or what might be expected *in vivo*. Submerged conditions can decrease ACE2 expression in human respiratory epithelial cells, thereby decreasing infection with SARS-CoV-2 virus (23); however, submerged conditions have been used successfully for SARS-CoV-2 pseudovirus transductions in other cell lines (8, 11) and for SARS-CoV-2 infection of respiratory cell lines (7). Nevertheless, use of an air-liquid interface format to enable more complete differentiation of respiratory epithelial cells might more accurately reflect the susceptibility of equine airways to SARS-CoV-2 infection. We did not examine the extent to which eACE2 is expressed by other regions of the equine respiratory tract. Ciliated cells have higher ACE2 expression and tropism for SARS-CoV-2 infection in humans and other species (24-26), but this has not yet been determined for horses. Expression of ACE2 varies among regions of the respiratory tract of humans and several domestic animal species, with greatest levels apically in the nasopharynx, decreasing levels in the conducting airways and lower airways, and then higher expression in alveolar cells (27-28). This pattern of ACE2 expression appears to be a limiting factor for susceptibility to infection *in vitro* and *in vivo* (27-30). Differences in expression of eACE2 among regions of the equine airway have not been well characterized. Expression of eACE2 in the alveoli and bronchi was not detected by immunohistochemistry in a single horse (26). However, ACE2 mRNA expression was detected in equine tracheal and lung tissue lysates (28). We have detected ACE2 expression using immunohistochemistry in equine alveoli and bronchi (unpublished data). Further investigation of variation in eACE2 expression and susceptibility to SARS-CoV-2 among epithelial cells of the equine airway is warranted.

Despite these limitations and the absence of evidence of SARS-CoV-2 infecting horses either experimentally (10) or naturally (5, 6, 18, 31), we demonstrated that EBECs are susceptible to infection with the SARS-CoV-2 pseudovirus. These findings suggest that SARS-CoV-2 can infect equine respiratory tract cells. If horses are susceptible to subclinical aerosol infection due to close contact with humans, they might serve as a potential reservoir for viral transmission to humans and for selection of viral mutations that render some strains more infective to horses (and possibly other animals). Horses in pasture might be exposed to wildlife infected with SARS-CoV-2, such as white-tailed deer that might have a relatively high prevalence of infection with SARS-CoV-2 (2). Serological and virological monitoring of horses in contact with persons shedding SARS-CoV-2 is warranted in light of seroconversion of a horse exposed to a person infected with SARS-CoV-2 (6), and contact with horses by humans infected with SARS-CoV-2 should be avoided due to potential zoonotic transmission. Although it is improbable at this time that horses represent a reservoir of SARS-CoV-2, further investigation of the susceptibility of horses to emerging strains of SARS-CoV-2 infection is warranted.

## MATERIALS AND METHODS

### Ethics statement

All methods were performed in accordance with relevant guidelines and regulations for animal use and for laboratory practices including environmental health, occupational safety, and biosafety (Texas A&M University Infectious Biohazard Committee IBC# 2017-105). Bronchial epithelial cells were harvested from horses euthanized for humane reasons unrelated to disease of the respiratory tract in accordance with practices approved by the Texas A&M University Institutional Animal Care and Use Committee.

### eACE2 protein sequence modelling

Angiotensin-converting enzyme 2 (ACE2) protein sequences for mammalian and avian species, selected based on likely human contact and exposure (*i*.*e*., pets and livestock) or wild animals documented as susceptible to infection (*e*.*g*., bats and whitetail deer), using a scheme adapted from a previous report (14) and updated to include horses, were retrieved from the National Center for Biotechnology Information Protein Database (NCBI, https://www.ncbi.nlm.nih.gov/, accession data in Supplementary Table 1). Sequence alignment was performed using the Clustal Omega plugin (http://www.clustal.org/omega) in Geneious Prime (Version 2021.2.2; https://www.geneious.com/) with default settings. Overall sequence homology to huACE2 (NCBI accession, #BDH16358.1), and specific amino acid residues identified as critical for interaction between the huACE2 protein and the receptor binding domain of SARS-CoV-2 (adapted from the work of Gultom and collaborators (14)) were compared amongst these species. Next, the 3-dimensional protein structure of eACE2 at its predicted interaction site with SARS-CoV-2 spike protein receptor binding domain (RBD) protein structure was modelled using the SWISS-MODEL web service (https://swissmodel.expasy.org). Its similarity to huACE2 was modelled using PDB structure 6m17 as a template.

### Culture of EBECs

EBECs were harvested postmortem from 5 horses with no history of respiratory disease. Primary cultures were established as previously described (15, 16). The lungs were removed en bloc within 1 hour of euthanasia, infused with 1 liter of ice-cold Hank’s buffered saline solution (HBSS) with penicillin (200 U/mL), streptomycin (200 µg/mL), and amphotericin B (2.5 µg/mL) into the trachea, and transported to the laboratory on ice. Lung parenchyma was bluntly removed to isolate primary and secondary bronchi, which were then sectioned in roughly 5-cm tubular segments and submerged for 30 minutes in ice-cold HBSS with penicillin (200 U/mL), streptomycin (200 µg/mL), and amphotericin B (2.5 µg/mL). The bronchial epithelium was sharply dissected from the underlying submucosa and then minced into approximately 1-mm^2^ segments and divided into batches of roughly 500 mg minced tissue. Each batch of tissue was placed into a Petri dish with 12 mL of 0.25% trypsin – 0.6 mM EDTA solution and incubated at 37°C for 2 hr in 5% CO_2_ under gentle agitation. Digestion was stopped by adding 5 mL ice-cold 20% fetal bovine serum (FBS) in HBSS. The suspension was filtered through sterile double-layered gauze, then a sterile cell strainer (pore size 40 µm; Greiner Bio-one Gmbh, Kremsmünster, Austria). The cellular suspension was centrifuged at 250 × g for 10 minutes at 4°C, the supernatant was discarded, and cells were resuspended in 12 mL warm supplemented airway culture media (Airway Epithelial Cell Growth Basal Media, Promocell; supplemented with Growth Medium SupplementPack [#C-39160, Promocell], 10% FBS, penicillin [200 U/mL], streptomycin [200 µg/mL], and amphotericin B [2.5 µg/mL]). This suspension was placed into a treated T75 cell culture flask and incubated for 30 minutes at 37°C in a humidified, 5% CO_2_ environment. The media was gently removed from the T75 flasks, allowing adhered fibroblasts to remain within the flask, and the cellular suspension containing EBECs was centrifuged at 200 × g for 10 minutes at 4°C. The supernatant was discarded and cells were resuspended in 1 mL of supplemented airway culture media. Cells were counted using a cell counter (CellometerAuto T4, Nexelom Bioscience, Lawrence, MA) with trypan blue for viability assessment, and plated at a concentration of 2.5 × 10^5^ live cells/ml on a collagen-coated tissue-treated 24-well plate. Media was changed at 24 hours, and then every 2 days afterward.

### Culture of HBECs

Human bronchial epithelial cells (HBECs, #C-12640, Promocell) were acquired as a cryopreserved positive control cell line. Primary cultures were established by thawing the cryovial in a 37°C water bath for 2 minutes and resuspending cells into warmed airway culture media (Airway Epithelial Cell Growth Basal Media, #C-21260, Promocell; supplemented with Growth Medium SupplementPack, #C-39160, Promocell), and plating at a density of 1.5 × 10^3^ cells/cm^2^ into a T25 flask. Media was changed at 24 hours, and then every 2 days afterward. Subsequent passages were plated into 24-well plates in parallel with EBECs to serve as a positive control primary cell line.

### Detection of ACE2 expression via western blot

EBECs were lysed with cold radioimmunoprecipitation assay (RIPA) lysis buffer and protease inhibitor (#P8340, Sigma Aldrich), then separated on 12% SDS-PAGE gels. Membranes were transferred to polyvinylidene difluoride (PVDF) membranes, blocked with 5% dried nonfat milk in 1% Tween-20 and phosphate-buffered saline (PBS), and probed with a polyclonal rabbit anti-huACE2 antibody targeting the C-terminus (1:1,000 dilution; #3217, ProSci) and a peroxidase-conjugated goat anti-rabbit secondary antibody (1:6,000 dilution; #711-035-152, Jackson ImmunoResearch Laboratories). Lysate of human lung adenocarcinoma A549 cells (#CCL-185, ATCC) was used as a positive control. For loading control, a separate aliquot of each sample during each immunoblot was probed with a monoclonal mouse anti-human GAPDH antibody (1:500 dilution; #43700, Invitrogen) and a peroxidase-conjugated goat anti-mouse secondary antibody (1:5,000 dilution; #6789, Abcam). Protein bands were developed using Radiance Plus® Femtogram HRP Substrate (Azure Biosystems) and visualized using the Bio-Rad Chemidoc Touch imaging system.

### Detection of ACE2 expression via flow cytometry

EBECs and HBECs were washed in PBS, lifted with Accutase (#07920, StemCell Technologies), counted, and divided into individual aliquots containing 1 × 10^6^ cells in 100 μL PBS containing 2% FBS. Cells were incubated with affinity-purified polyclonal goat anti-huACE2 antibody recognizing the extracellular domain (1:40 dilution, #AF933, R&D Systems) for 30 minutes, washed twice, and resuspended with 300 μL of PBS containing 2% FBS. Cells were then stained with a secondary rabbit anti-goat Alexa Fluor 555 antibody (1:400 dilution, #A32732, Invitrogen) for 30 minutes at room temperature, washed twice and resuspended in 100 µl PBS containing 2% FBS. The cells were then fixed with 2% paraformaldehyde for 30 minutes. The ACE2 Alexa Fluor 555 labeled fixed EBECs and HBECs were run on a Luminex/Amnis Image Stream X Mark II imaging flow cytometer to measure the amount of ACE2 expressed in the cells. Settings for cytometry using the INSPIRE (Luminex/Amnis) acquisition software were as follows: 1) the 40× objective was used to collect the images and light from the cells interrogated by the laser; 2) the 488-nm laser was set at 100 mW of power; 3) the side-scatter laser (785 nm) was set at 2 mW of power; 4) the core diameter was set at 10 μm; and, 5) the fluid velocity was set at 132 mm/sec. At least 10,000 events were collected for each sample. IDEAS software (version 6.3, Luminex/Amnis) was used to analyze the data collected from the Image Stream. The gating strategy (Supplemental Figure 1) was set up with singlets being selected with the y-axis being the Aspect Ratio of Channel 1 bright-field and the x-axis being the area of Channel 1 bright-field. Cells with an aspect ratio >0.6 were selected as single cells. The single cells were then gated for in-focus cells by the cells with the greater gradient RMS values in Channel 1 bright-field. The single, focused cells were then observed for Alexa Fluor 555-positive cells by plotting intensity of channel 2 (FITC channel) on the y-axis and the intensity of channel 3 (Alexa Fluor 555) on the x-axis. The ACE2 Alexa Fluor 555-positive cells were gated as the population shifted toward the Alexa Fluor 555 direction away from the autofluorescence background of the cells. The proportions of ACE2-positive EBECs and HBECs were compared using chi-squared test.

### Detection of ACE2 expression using immunofluorescence

EBECs were plated at a density of 250,000 cells/chamber in 0.1% collagen-coated ibiTreat µ-Slide 2-well chambered coverslips (Ibidi) and allowed to grow over 72 hours, changing media daily. Cells were then washed with PBS, fixed with PBS buffered 4% paraformaldehyde solution, and blocked with 10% goat serum (Sigma Aldrich) in PBS for 2 hours. Slides were then incubated with primary polyclonal rabbit anti-huACE2 antibody (#3217, ProSci) at 1:240 dilution in 1% BSA overnight. After incubation, cells were washed 3 times with PBS and incubated with secondary goat anti-rabbit Alexa Fluor 488 antibody (#A11008, Invitrogen) at 1:500 dilution in 1% BSA for 2 hours, followed by 2 µg/mL of Hoechst 33342 (#H3570, Invitrogen) in 1% BSA for 1 minute. After incubation, cells were washed 3 times with PBS, treated with 1 drop of Prolong Gold Antifade Reagent (#P36930, Invitrogen), and covered with a micro-coverslip. An additional set of fixed EBECs were permeabilized with 0.1 % Triton-100X in PBS for 12 minutes at room temperature, washed 3 times with PBS and blocked with 10% goat serum in PBS for 2 hours. To confirm epithelial cell-type, permeabilized EBECs were incubated with murine AE1/AE3 pan-cytokeratin antibody (#CM011A, Biocare Medical) at 1:100 dilution in 1% BSA in PBS at 4°C overnight, followed by a secondary goat anti-mouse Alexa Fluor 594 antibody (1:500 dilution, #8890, Cell Signaling Technologies) for 2 hours at 4°C prior to applying a coverslip as previously described. Slides were imaged immediately following staining using a Zeiss 780 confocal microscope with laser excitation of 405 nm and 594 nm, and emission was collected using wavelengths of 410 to 480 nm and 596 to 700 nm, respectively. In a subsequent experiment, EBECs and HBECs were plated at the same density and under similar conditions, and processed as previously described, with the exception of secondary staining with secondary goat anti-rabbit Alexa Fluor 555 antibody (#A32732, Invitrogen) at 1:500 dilution in 1% BSA for 2 hours, followed by 2 µg/mL of Hoechst 33342 in 1% BSA for 1 minute.

### Pseudovirus transduction of EBECs

A commercially acquired lentiviral construct pseudotyped with the spike (S) protein of SARS-CoV-2 (GenBank accession no. NC_045512.2) and expressing enhanced green fluorescence protein (eGFP) (vector VB900088-2229upx, VectorBuilder) was used as a reporter. Transduction using approximately 132,000 transduction units (TU) of pseudovirus and 1 µg/mL polybrene (VectorBuilder), was performed in cultured EBECs derived from 3 horses; HBECs obtained commercially were included as positive controls. Transduced and non-transduced EBECs and HBECs were examined after 96 hours by fluorescence microscopy for evidence of green fluorescence. Images were obtained with an Evos FL microscope, using the EVOS GFP light cube (ThermoFisher Scientific), with excitation of 482/25 nm and emission of 524/24 nm. RNA was extracted from bronchial cells treated with TRIzol reagent (ThermoFisher), using a commercial kit (Direct-zol DNA/RNA Miniprep Kit) according to the manufacturer’s instructions. DNA was reverse-transcribed from 0.5 µg of RNA, using a commercial kit (RT^2^ First Strand Kit, Qiagen). Quantification of transduction of EBECs and HBECs with the SARS-CoV-2 pseudovirus was determined by real-time qPCR (rt-PCR) for gene expression of the enhanced green fluorescence protein gene (*egfp*); expression of the β-2 microglobulin gene (*B2M*) was used as a comparative reference. A no-template preparation (negative control) and a plasmid encoding *egfp* (positive control, pClneoEGFP human RASSF6b, Addgene plasmid #37021), along with non-transduced EBECs and HBECs were included. The rt-PCR reaction used 1 μL containing 50 ng of cDNA, added to 5 μL of a commercial master mix (TaqMan Fast Advanced, Applied Biosystems), 0.5 μL of a primer/probe premix against either eGFP, equine or human B2M (for YFP∼YFP (eGFP): FAM-MGB, assay# Mr04097229_mr; for equine B2M: FAM-MGB, assay# Ec03468699_m1; for human B2M: FAM-MGB, assay# Hs00187842_m1; TaqMan Gene Expression Assays, Applied Biosystems), and 3.5 μL of buffered nuclease-free water. Each condition was repeated in triplicate, and each sample from the 3 experiments was tested in triplicate by qPCR. Samples were processed using commercial software for rt-PCR (QuantStudio Real-Time PCR Software v1.3, Applied Biosystems). Ct values of *egfp* expression were compared among treatments using generalized linear modelling, with post hoc comparisons made using the method of Tukey. The ΔCt values (i.e., Ct[*egfp*] – Ct[*B2M*]) were determined for EBECs and HBECs and compared using the Wilcoxon rank sum test.

## Acknowledgements

The project was funded by the Link Equine Research Endowment, Texas A&M University, and the Department of Large Animal Clinical Sciences, School of Veterinary Medicine & Biomedical Sciences, Texas A&M University. The authors acknowledge the assistance of the Image Analysis Laboratory (RRID: SCR_022479) and Flow Cytometry Facility (RRID: SCR_022169), School of Veterinary Medicine & Biomedical Sciences, Texas A&M University. The authors thank Mrs. Jocelyne Bray for her technical assistance.

## Figure Legends

**Supplementary Table 1.** ACE2 Protein Sequence Accession Data. ACE2 protein sequences and accession numbers used for alignment.

Sequence Accession Data
| Common name | Species name | Accession Number |
| --- | --- | --- |
| American Mink | *Neogale vison* | XP_044091953.1 |
| Bactrian Camel | *Camelus bactrianus* | XP_010966303.1 |
| Bat: Carolia | *Carollia perspicillata* | QJF77828.1 |
| Bat: Rhinolophus | *Rhinolophus sinicus* | AGZ48803.1 |
| Bat: Sturnira | *Sturnira lilium* | QWM88991.1 |
| Cattle: Indicus | *Bos indicus x Bos taurus* | XP_027389727.1 |
| Cattle: Taurus | *Bos taurus* | XP_005228485.1 |
| Chinese Pangolin | *Manus pentadactyla* | XP_036768816.1 |
| Domestic Cat | *Felis catus* | XP_044906240.1 |
| Domestic Chicken | *Gallus gallus* | XP_416822.3 |
| Domestic Dog | *Canis lupus familiaris* | QLF98519.1 |
| Domestic Goat* | *Capra hircus* | XP_005701129.2 |
| Domestic Rabbit | *Oryctolagus cuniculus* | QHX39726.1 |
| Donkey | *Equus asinus* | XP_044620396.1 |
| Ferret | *Mustela putorius furo* | XP_004758942.1 |
| Horse | *Equus caballus* | XP_001490241.1 |
| Human | *Homo sapiens* | BDH16358.1 |
| Llama | *Lama glama* | QWM88990.1 |
| Macaque | *Macaca mulatta* | XP_028697658.1 |
| Sheep | *Ovis aries* | XP_042098228.1 |
| White-Tailed Deer | *Odocoileus virginianus texanus* | XP_020768965.1 |
| Wild Boar | *Sus scrofa* | XP_020935033.1 |

**Supplementary Figure 1.**
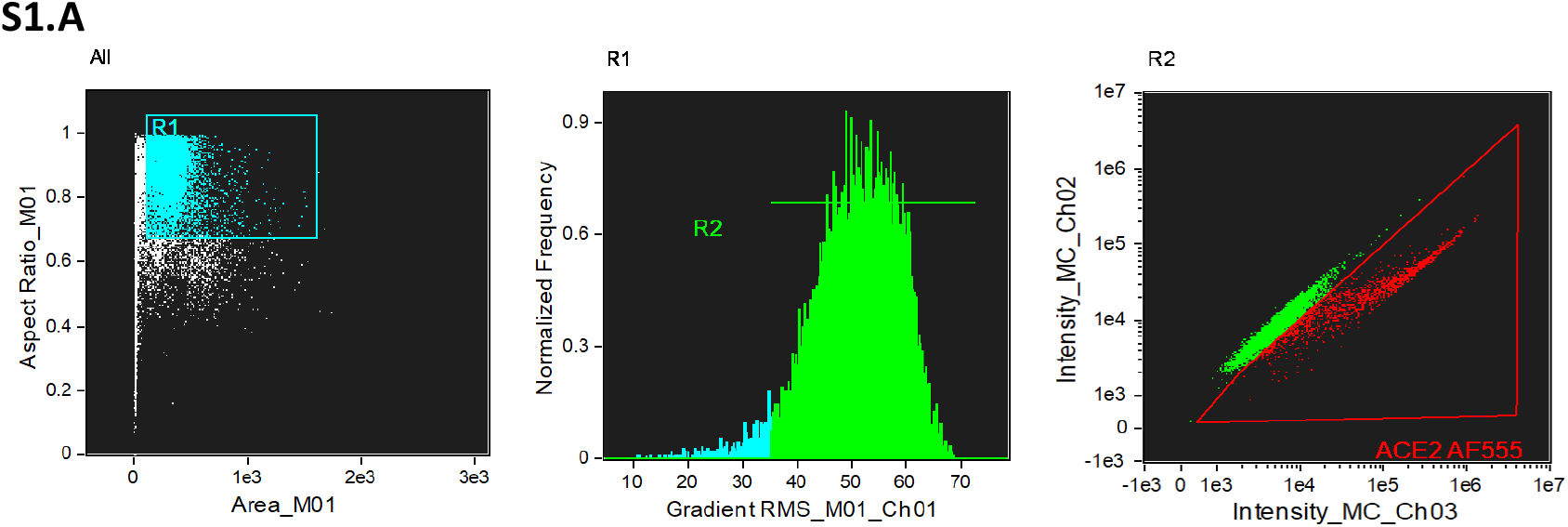
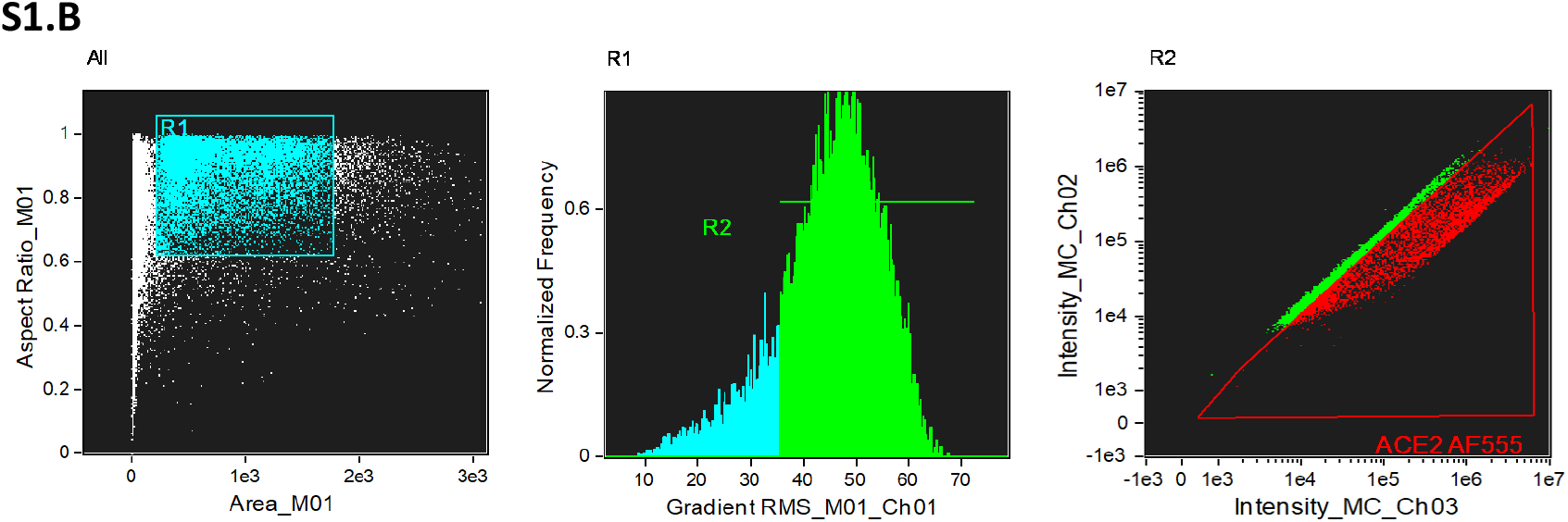
A EBEC ACE2 AF 555 gating strategy B. HBEC ACE2 AF 555 Gating Strategy. Gating strategy used for imaging flow cytometry. **(A)** gating strategy of cultured EBECs expressing ACE2. The left scatterplot shows the aspect ratio of Channel 1 (Brightfield) on the y-axis and the area of Channel 1 (Brightfield) on the x axis and the population of single EBEC cells were selected with the R1 gate. Cells with an aspect ratio >0.6 were selected as single cells. The center histogram depicts the gradient RMS values on the x-axis and the EBECs that were in focus were selected with gate R2. In focus EBECs exhibited a gradient RMS value of >35. The right scatterplot shows the intensity of Channel 3 (AF555 channel) on the x-axis and the intensity of Channel 2 (autofluorescence channel) on the y-axis and the EBECs expressing ACE2 AF555 were selected with gate ACE2 AF555. The ACE2 Alexa Fluor 555-positive cells were gated as the population shifted toward the Alexa Fluor 555 direction away from the autofluorescence background of the cells. **B)** gating strategy of cultured HBECs expressing ACE2. The same gating strategy was used for the HBECs as was used for the EBECs explained in panel A.

